# The two formyl peptide receptors differently regulate GPR84-mediated neutrophil NADPH-oxidase activity

**DOI:** 10.1101/2020.11.13.381582

**Authors:** Jonas Mårtensson, Martina Sundqvist, Asmita Manandhar, Loukas Ieremias, Linjie Zhang, Trond Ulven, Xin Xie, Lena Björkman, Huamei Forsman

## Abstract

Neutrophils express many G protein-coupled receptors (GPCRs) including the two formyl peptide receptors (FPR1 and FPR2) and the medium chain fatty acid receptor GPR84. The FPRs are known to define a hierarchy among neutrophil GPCRs, i.e., the GPCR-mediated response can be either suppressed or amplified by signals generated by FPRs. In this study, we investigated the position of GPR84 in the FPR-defined hierarchy regarding the activation of neutrophil NADPH-oxidase, an enzyme system designed to generate reactive oxygen species (ROS). When naïve neutrophils are activated by GPR84 agonists a modest ROS release was induced. However, vast amounts of ROS production was induced by these GPR84 agonists in FPR2-desensitized neutrophils, and the response is inhibited not only by a GPR84 antagonist but also by an FPR2 specific antagonist. This suggests that the amplified GPR84 agonist response is achieved through a reactivation of the desensitized FPR2. In addition, the GPR84-mediated FPR2 reactivation was independent of β-arrestin recruitment and sensitive to a protein phosphatase inhibitor. In contrast, the modest ROS production induced by GPR84 agonists was primarily suppressed in FPR1-desensitized neutrophils through hierarchical desensitization of GPR84 by FPR1 generated signals.

In summary, our data show that FPRs control the NADPH-oxidase activity mediated through GPR84 in human neutrophils. While an amplified ROS generation is achieved by GPR84 agonists through reactivation of desensitized FPR2, FPR1 heterologously desensitizes GPR84 and by that suppresses the release of ROS induced by GPR84 agonists.

## INTRODUCTION

Neutrophils, major effector cells in innate immunity, constitute a vital part of the defense against invading microbes and other protective inflammatory processes, but can also mount pathological actions leading to tissue damage (1–3). The multifaceted functions of neutrophils, including directional migration, secretion of granule constituents and activation of the reactive oxygen species (ROS) generating nicotine adenine dinucleotide phosphate (NADPH)-oxidase, are regulated and fine-tuned by different receptors belonging to the family of G protein-coupled receptors (GPCRs) (4, 5). Neutrophils express a number of GPCRs of importance for initiation as well as resolution of inflammation, including the two formyl peptide recognizing receptors (FPR1 and FPR2), the ones that recognize platelet activating factor (PAFR), interleukin 8 (CXCR1/2), and the free fatty acids (FFARs) (4, 6, 7). ROS generated by the NADPH-oxidase play roles not only in bacterial killing and tissue destructing activities but also in cell signaling and immune regulation (8–11). Thus, it is of outmost importance that activation of the oxidase is a well-regulated process; an efficient inflammatory response should be achieved without causing any unnecessary collateral tissue damage.

The neutrophil response triggered by GPCRs relies not only on the specific agonists that activate their respective receptor, but also on a complex receptor down-stream signaling network and different hierarchical receptor cross-talk mechanisms that regulate receptor activities and fine-tune the functions of neutrophils (7). The complexity can be illustrated by a defined position of receptor hierarchical cross-talk between FPRs and other GPCRs. For example, on one hand, signals generated down-stream of the FPRs heterologously desensitize CXCR1/2, and totally abrogate IL-8 induced neutrophil functions (12–15). On the other hand, FPR agonists amplify the response induced by PAF and ATP through a receptor cross-talk mechanism with the PAFR and the P2Y2R (receptor for ATP), respectively (16–18).

GPR84 senses the medium-chain fatty acids (MCFAs) such as lauric acid and capric acid that are available through food intake (19). Based on the facts that GPR84 recognizes MCFAs and is highly expressed by neutrophils, this receptor has been suggested to constitute a molecular link between metabolism and inflammation (19–21). The precise activation mechanism(s) and the physiological role of GPR84 remains still largely unknown but the identification of several potent GPR84-selective agonists as well as antagonists over the last years has greatly increased our understanding of this receptor (see the two recent reviews (22, 23)). The availability of these powerful molecular tool compounds makes it possible to perform mechanistical studies focusing on GPR84 signaling and regulation.

In this study, we have used new GPR84-related molecular tools to determine the regulatory effects of FPRs on GPR84 triggered activation of neutrophils. The data obtained clearly shows that FPR1 as well as FPR2 affects GPR84-mediated neutrophil function. Our results add new knowledge regarding the complexity of neutrophil GPCR communication hierarchy demonstrated by that the two very closely related FPRs regulate the function of GPR84 differently. Whereas FPR1 primarily suppressed the GPR84 induced production of ROS, a suppression suggested to be induced by heterologous GPR84 desensitization, FPR2 positively augmented the response induced by GPR84 specific agonists. The basis for the increased ROS production induced by GPR84 agonists in FPR2-desensitized cells is disclosed to involve a novel receptor cross-talk mechanism by which signals generated by GPR84 reactivate desensitized FPR2.

## MATERIALS AND METHODS

### Ethics statement

This study was conducted using human blood neutrophils from buffy coats obtained from the blood bank at Sahlgrenska University Hospital, Gothenburg Sweden. The Swedish legislation section code 4§ 3p SFS 2003:460 (Lag om etikprövning av forskning som avser människor), states that no ethical approval is needed for research on buffy coats, as they were provided anonymously and thereby cannot be linked to a specific donor.

### Chemicals

Dextran and Ficoll-Paque were obtained from GE-Healthcare Bio-Science (Waukesha, WI, USA). Horseradish peroxidase (HRP) was obtained from Roche Diagnostics (Mannheim, Germany). Isoluminol, dimethyl sulfoxide (DMSO) and the formylated tripeptide *N*-formyl-Met-Leu-Phe (fMLF) were obtained from Sigma (Sigma Chemical Co., St. Louis, MO, USA). Cyclosporine H was kindly provided by Novartis Pharma (Basel, Switzerland). The FPR2 pepducin F2Pal10, the PIP2-binding peptide PBP10, the hexapeptide WKYMVM, the mitochondrial cryptic peptide (mitocryptide) NADH dehydrogenase subunit 4 (MCT-ND4) and the formylated tetrapeptide *N*-formyl-Met-Ile-Phe-Leu (fMIFL) were synthesized and HPLC-purified by TAG Copenhagen A/S (Copenhagen, Denmark). Calyculin A was purchased from Nordic Biosite (Täby, Sweden) and the Src family kinase inhibitor PP2 were from Calbiochem (La Jolla, CA, USA). PAF was obtained from Avanti Polar Lipids (Alabaster, AL, USA) and ZQ16 (2-(hexylthio)pyrimidine-4,6-diol) was from Tocris Bioscience (Bristol, UK). The earlier described GPR84 specific agonists Cpd51 (6-nonylpyridine-2,4-diol) and DL175 were synthesized as described (24, 25). The antagonist GLPG1205 was kindly provided by Galapagos NV (Mechelen, Belgium). All peptides were dissolved in DMSO and stored at −80°C until use. Subsequent dilutions of all peptides were made in Krebs-Ringer phosphate buffer (KRG, pH 7.3; 120 mM NaCl, 5 mM KCl, 1.7 mM KH_2_PO_4_, 8.3 mM NaH_2_PO_4_ and 10 mM glucose) supplemented with Ca^2+^ (1 mM) and Mg^2+^ (1.5 mM).

### Isolation of human neutrophils

Human peripheral blood neutrophils were isolated from buffy coats from healthy blood donors using dextran sedimentation and Ficoll-Paque gradient centrifugation as described (26, 27). The remaining erythrocytes were disrupted by hypotonic lysis and the neutrophils were washed twice, and resuspended in KRG and stored on melting ice until use. This isolation process permits cells to be purified with minimal granule mobilization.

### Neutrophil NADPH-oxidase activity

The NADPH-oxidase activity was determined using isoluminol-enhanced chemiluminescence (CL) (28). The CL activity was measured in a six-channel Biolumat LB 9505 (Berthold Co., Wildbad, Germany), using disposable 4-ml polypropylene tubes with a 900-microliter reaction mixture containing 10^5^ neutrophils, isoluminol (2 x 10^-5^ M) and HRP (2 Units/ml). The tubes were equilibrated in the Biolumat for five minutes at 37°C, after which the stimulus (100 μl) was added and the light emission was recorded continuously over time. The light emission/superoxide anion production is expressed as Mega counts per minute (Mcpm).

When receptor desensitized cells were studied, naïve (non-desensitized) cells were first stimulated with a receptor-specific agonist and when the release of superoxide had declined, a second stimulation was performed. When experiments were performed with antagonists, these were added to the neutrophils in the CL reaction mixture one minute before the second stimulation.

### GPR84-mediated cyclic adenosine monophosphate (cAMP) assay

HEK293 cells stably expressing GPR84 were harvested and re-suspended DMEM containing 500 μM IBMX (an inhibitor of cyclic nucleotide phosphodiesterases) at a density of 4 × 10^5^ cells/ml. Cells were then plated onto 384-well assay plate at 2000 cells/5 μl/well. Another 5 μl buffer containing compounds at various concentrations were added to the cells. After incubation at room temperature for 30 minutes, 5 μl DMEM containing 15 μM forskolin was added, and the incubation was continued for another 30 minutes. Intracellular cAMP measurement was carried with a LANCE Ultra cAMP kit (PerkinElmer, Cat No: TRF0264, Waltham, USA) using an EnVision multiplate reader (PerkinElmer, Waltham, USA) according to the manufacturer’s instructions.

### GPR84-mediated β-arrestin2 recruitment

The recruitment of β-arrestin2 to GPR84 was measured using the Promega NanoBiT Protein-Protein Interaction System (29). In brief, HEK293 cells seeded at 4 × 10^5^ cells/well on 96-well plates in DMEM media were transfected with plasmids encoding Smbit-β-arrestin2 and GPR84-LgBit. Twenty-four hours later, the cells media were replaced with 40 μl fresh culture medium without FBS. Thereafter 10 μl Nano-Glo Live Cell reagent was added according to the manufacturer’s protocol (Promega, Cat No: N2011, Madison, USA), and the cells were incubated in a 37°C, 5% CO2 incubator for five minutes before another 25 μl culture media with various concentrations of compounds were added to the cells. After incubation at room temperature for 15 minutes, bioluminescence was measured with an EnVision multiplate reader 2104 (PerkinElmer, Waltham, USA).

### Data analysis

Data processing and analysis was performed using GraphPad Prism 8.0.0 (Graphpad Software, San Diego, CA, USA). Curve fitting was performed by non-linear regression using the sigmoidal dose-response equation (variable-slope). For statistical analysis, either a paired Student’s *t*-test or a repeated measurement one-way ANOVA followed by Dunnett’s multiple comparison test was used as indicated in the figure legends. All statistical analysis was performed on raw data values and statistically significant differences are indicated by \**p* < 0.05 or \*\**p* < 0.01.

## RESULTS

### Naïve neutrophils activated by GPR84 specific agonists release reactive oxygen species (ROS)

Activation of neutrophils by many GPCR agonists triggers an assembly of the NADPH-oxidase and a release of ROS (4). Accordingly, when naïve human neutrophils were stimulated with the two GPR84 specific agonists ZQ16 and its structural analog Cpd51 ((24), structures shown in Fig 1A-B), the NADPH-oxidase was activated to produce ROS (Fig 1C). The poor ROS response mediated through GPR84 is in line with our earlier results (21). The ROS production profile mediated by the GPR84 agonists was very similar to that induced by the FPR2 specific agonistic pepducin F2Pal10, i.e., the response was induced rapidly upon agonist stimulation and was terminated within three minutes after initiation (Fig 1C). However, the maximal GPR84 agonist-induced ROS production from naïve neutrophils was low in comparison to that induced by a standard concentration (500 nM) of F2Pal10 (Fig 1C-D). These data show that both FPR2 and GPR84 agonists trigger an activation of the neutrophil NADPH-oxidase, but that the magnitude of the GPR84-induced ROS response is less pronounced as compared to the response induced by the FPR-agonists.

**Figure 1.**
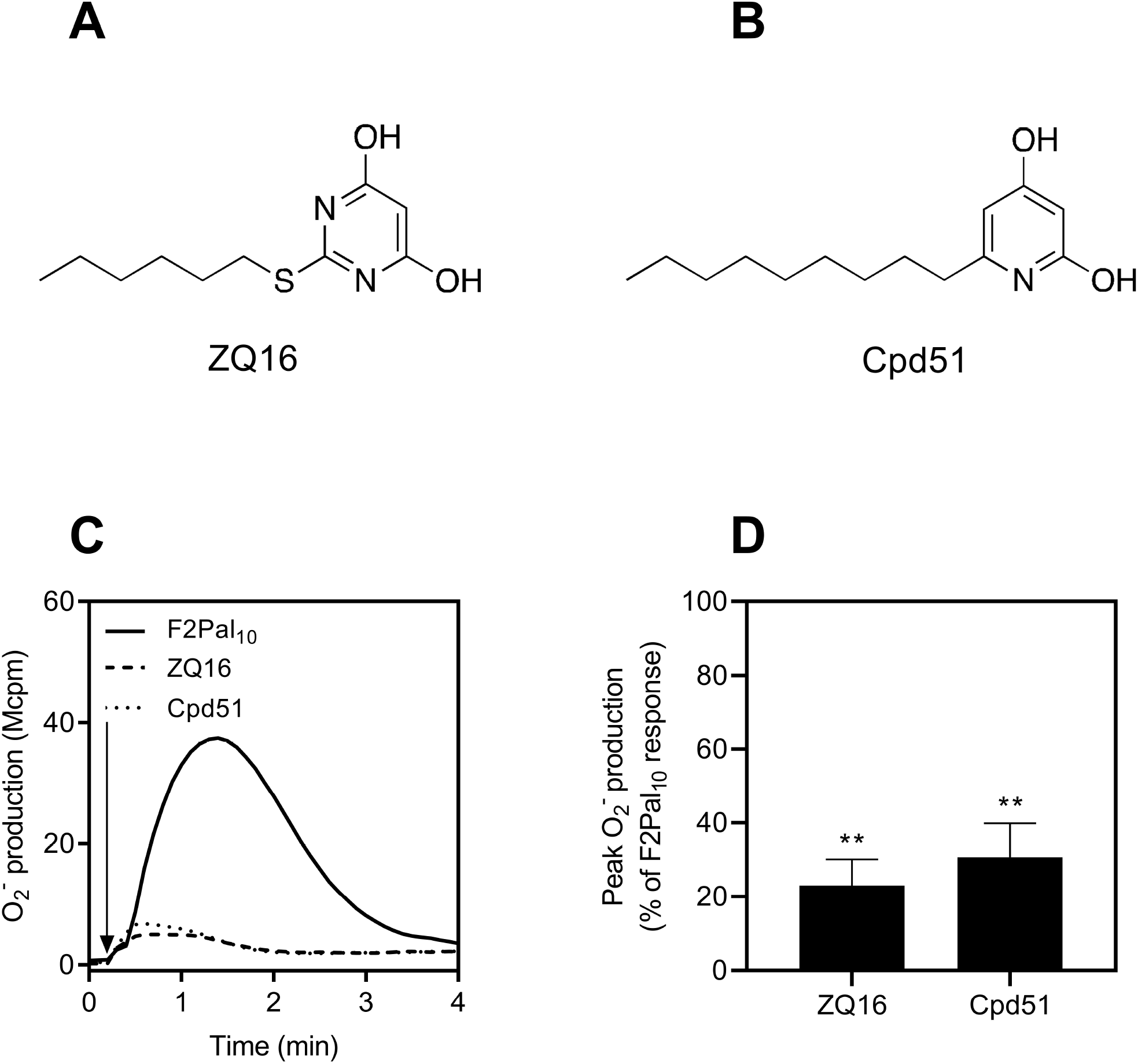
Chemical structures of GPR84 agonists and their ability to activate the neutrophil NADPH-oxidase. Chemical structures of the GPR84 agonists **A**) ZQ16 and **B**) the ZQ16 structural related analog Cpd51. **C-D**) Naïve neutrophils were stimulated with F2Pal_10_ (500 nM), ZQ16 (1 μM) or Cpd51 (1 nM) after which the release of superoxide was measured continuously over time. **C**) One representative trace of the superoxide anion release and **D**) A summary of the peak superoxide anion production presented in bar graph as % of the F2Pal_10_ response (mean + SEM, n = 4).

### The GPR84-mediated ROS release is largely amplified in FPR2-desensitized neutrophils

As shown, ROS production following activation of FPR2 is rapidly terminated (Fig 1C). When the activity has returned to basal levels, these neutrophils are non-responsive (desensitized) to additional activation with same agonist (shown with F2Pal_10_ in Fig 2A) as well as other FPR2 selective agonists (30). However, the level of ROS released from F2Pal_10_ desensitized neutrophils when triggered by ZQ16 was significantly increased when compared to the ZQ16 response from naïve neutrophils (Fig 2A-B). The level of the ZQ16-induced ROS production in FPR2-desensitized neutrophils was dependent on the F2Pal_10_ concentration used to desensitize the receptor, reaching a maximum level with ~ 250 nM F2Pal_10_ (Fig 2B). The enhanced level of ZQ16-induced ROS production in FPR2-desensitized neutrophils was not restricted to the use of F2Pal_10_ as the desensitizing agonist, as similar amplifications were obtained also when FPR2 was desensitized with the FPR2 agonists MCT-ND4 (a mitochondrial-derived formyl peptide; Fig 2C) and the hexapeptide WKYMVM (Fig 2D). Despite the fact that the highest concentrations of FPR2 agonists did not induce the highest degree of amplification of the ZQ16 response, no inhibitory effect on the ZQ16 response was obtained with increasing concentrations of either of the FPR2 agonists tested (Fig 2B-D). As such, F2Pal_10_ was used for further and more detailed characterization of the increased GPR84-mediated ROS response in FPR2-desensitized neutrophils.

**Figure 2.**
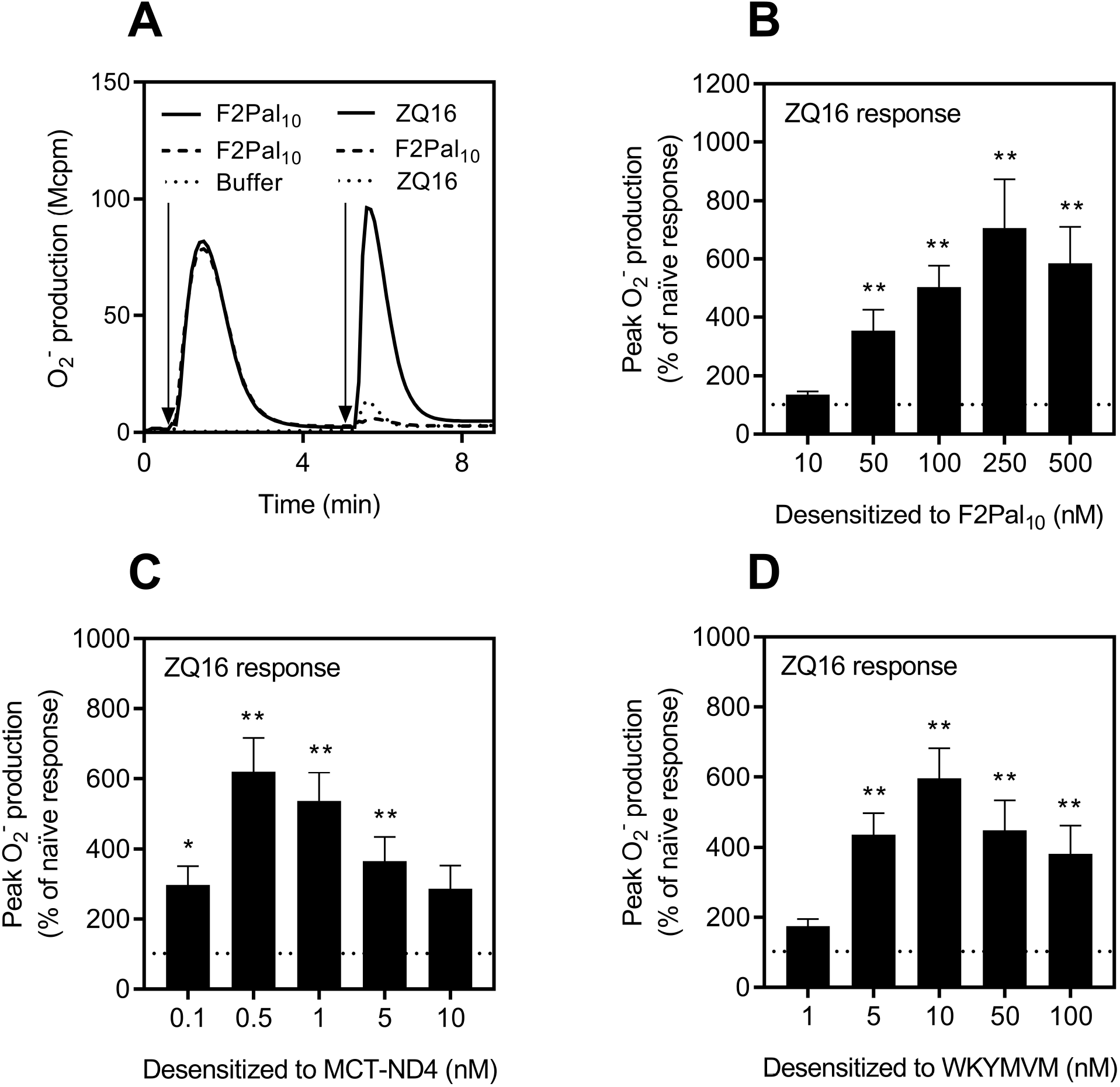
Enhanced reactivation induced by ZQ16 in FPR2-desensitized neutrophils. **A**) Naïve neutrophils were first activated with F2Pal_10_ (500 nM, solid and dashed lines) or buffer (dotted line) and the release of superoxide anions was measured continuously over time. After five min, when the F2Pal_10_ response had returned to basal levels, the neutrophils received a second stimulation with F2Pal_10_ (500 nM, dashed lines) or ZQ16 (1 μM, solid and dotted lines). One representative experiment out of five individual experiments for each agonist combination is shown. **B-D**) Summary of the second response induced by ZQ16 (1 μM) from FPR2 desensitized neutrophils pre-activated with different concentrations of **B**) F2Pal_10_ (n = 5), **C**) MCT-ND4 (n = 5) or **D**) WKYMVM (n = 5). The second peak superoxide response was determined and bar graphs show the percent (mean + SEM) of the naïve ZQ16 response (neutrophils that received buffer as the first stimulation) denoted as a response of 100% and indicated by the dotted line. Statistically significant differences in B-D were assessed using a repeated measurement one-way ANOVA followed by Dunnett’s multiple comparison test to the naïve ZQ16 response.

The response induced by the ZQ16 analog Cpd51 (24), was also significantly amplified in F2Pal_10_-desensitized neutrophils (Fig 3A), reaching a maximum level already at ~ 50 nM F2Pal_10_ (Fig 3B). Furthermore, the magnitude of the ROS production induced by both ZQ16 and Cpd51 in F2Pal_10_-desensitized neutrophils was similar to that induced by F2Pal_10_ in naïve neutrophils (Fig 2A and 3A). However, it should be noted that the GRP84 agonists, when compared to F2Pal_10_, were very poor ROS inducers in naïve cells (Fig 1D). In addition, when titrating the concentrations of the GPR84 agonists used to induce ROS in neutrophils desensitized with a fixed concentration of F2Pal_10_ a dose-dependent amplification of the response was revealed; EC50 value for Cpd51 ~ 0.2 nM (Fig 3C) and EC50 value for ZQ16 ~ 200 nM (Fig 3D). Thus, even though Cpd51 was ~ 1000 times more potent than ZQ16 at inducing NADPH-oxidase derived ROS-production in FPR2-desensitized, these data clearly demonstrate that FPR2 activation/desensitization transfers GPR84 agonists from poor to potent ROS activating agonists.

**Figure 3.**
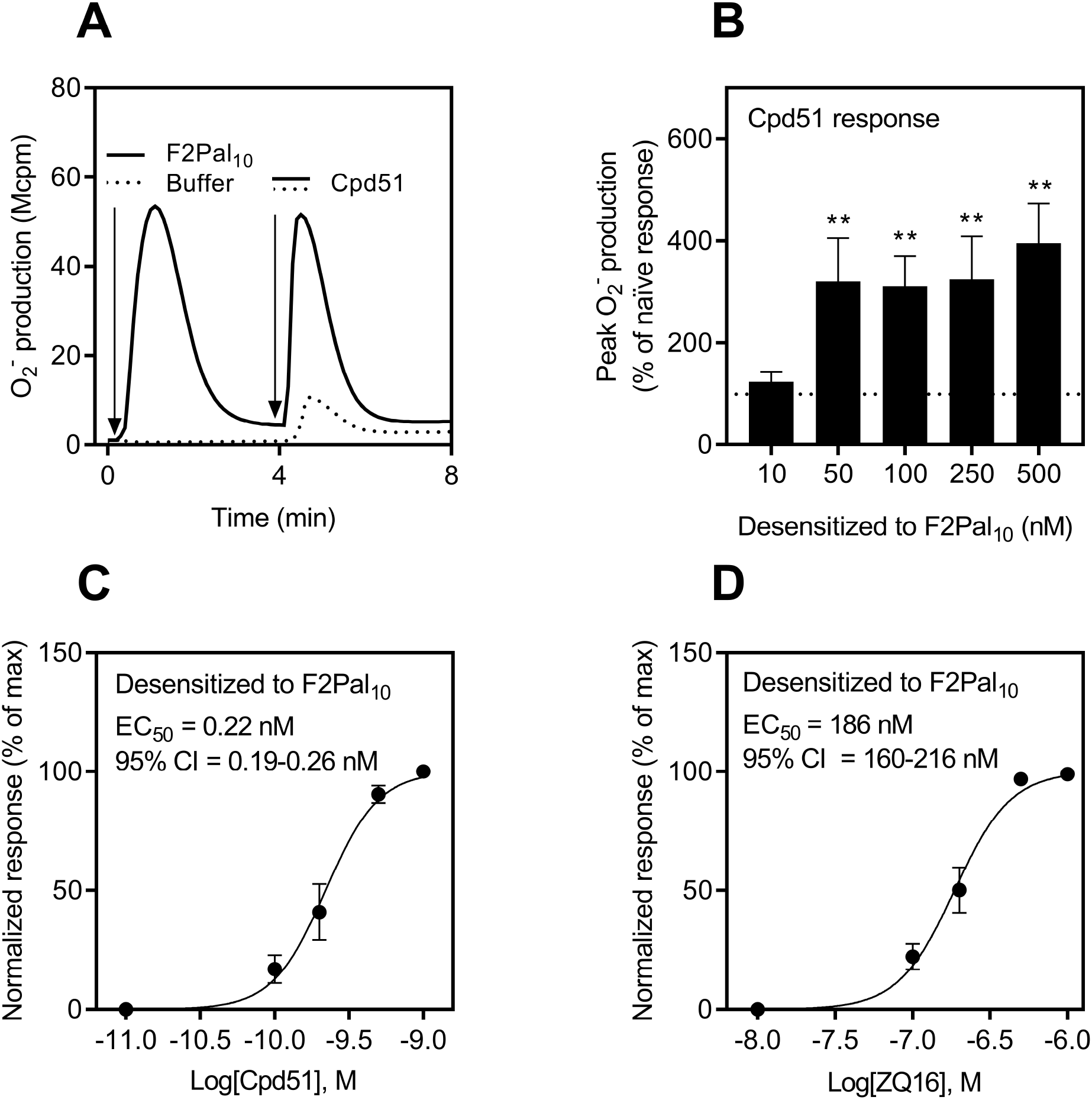
F2Pal_10_ transfers GPR84 agonists into potent NADPH-oxidase activators. **A**) Naïve neutrophils were first activated with F2Pal_10_ (500 nM, solid line) or buffer (dotted line) and the release of superoxide anions was measured continuously over time. When the F2Pal_10_ response had returned to basal levels, the neutrophils received a second stimulation with Cpd51 (1 nM, solid and dotted lines). One representative experiment out of four individual experiments for each agonist combination is shown. **B**) Summary of the second GPR84-agonist induced response from neutrophils pre-activated with different concentrations F2Pal_10_ (mean + SEM, n = 4). Statistically significant differences from the naïve Cpd51 response were assessed using one-way ANOVA followed by Dunnett’s multiple comparison test to the naïve Cpd51 response (denoted as a response of 100% and indicated by the dotted line). **C-D**) Dose-response curves of the second response induced by **C**) Cpd51 or **D**) ZQ16 from FPR2 desensitized cells pre-activated with a fixed concentration of F2Pal_10_ (500 nM). Data were normalized to the maximal response induced by the respective agonists. The EC50 values and 95% confidence interval (CI) were calculated based on the peak ROS production (mean + SEM, n = 3 independent experiments).

### GPR84 triggers a unidirectional reactivation of desensitized FPR2

To characterize the precise receptor involvement of the amplified GPR84 response in FPR2-desensitized neutrophils, we determined the effects of antagonists specific for FPR2 and GPR84, respectively. The receptor selectivity of the two antagonists PBP10 (selective for FPR2; (31)) and GLPG1205 (selective for GPR84; (32)) was confirmed using F2Pal_10_ and Cpd51 as neutrophil activating agonists. As expected, the FPR2 antagonist inhibited the response induced by F2Pal_10_ (Fig 4A) and was without effect on the response induced by Cpd51 (Fig 4B), whereas the GPR84 antagonist inhibited the Cpd51 response (Fig 4B) and was without effect on the F2Pal_10_ induced response (Fig 4A). Interestingly, the response induced by the GPR84 agonist ZQ16 in FPR2-desensitized neutrophils was significantly inhibited by PBP10; the FPR2 antagonist was added at a time point when the F2Pal_10_ induced response had been terminated (Fig 4C, D). The degree of inhibition by PBP10 was comparable to that induced by the GPR84 antagonist GLPG1205, and the inhibited response reached similar levels to that induced by ZQ16 in naïve neutrophils (Fig 4D). Very similar inhibitory patterns were observed with the two antagonists when ZQ16 was replaced by the other GPR84 agonist Cpd51 (Fig 4E). These data thus suggest that GPR84 can cross-talk with FPR2 and the GPR84-mediated ROS production in FPR2-desensitized neutrophils involves a reactivation of desensitized FPR2. In addition, the GPR84 antagonist was without effect when the order of the activating agonists was reversed. That is, the F2Pal_10_ response in GPR84-desensitized neutrophils was not affected by the GPR84 antagonist (Fig 4F). This strongly implies that the receptor cross-talk is unidirectional, i.e., GPR84 transduces signals leading to FPR2 reactivation, but not *vice versa*.

**Figure 4.**
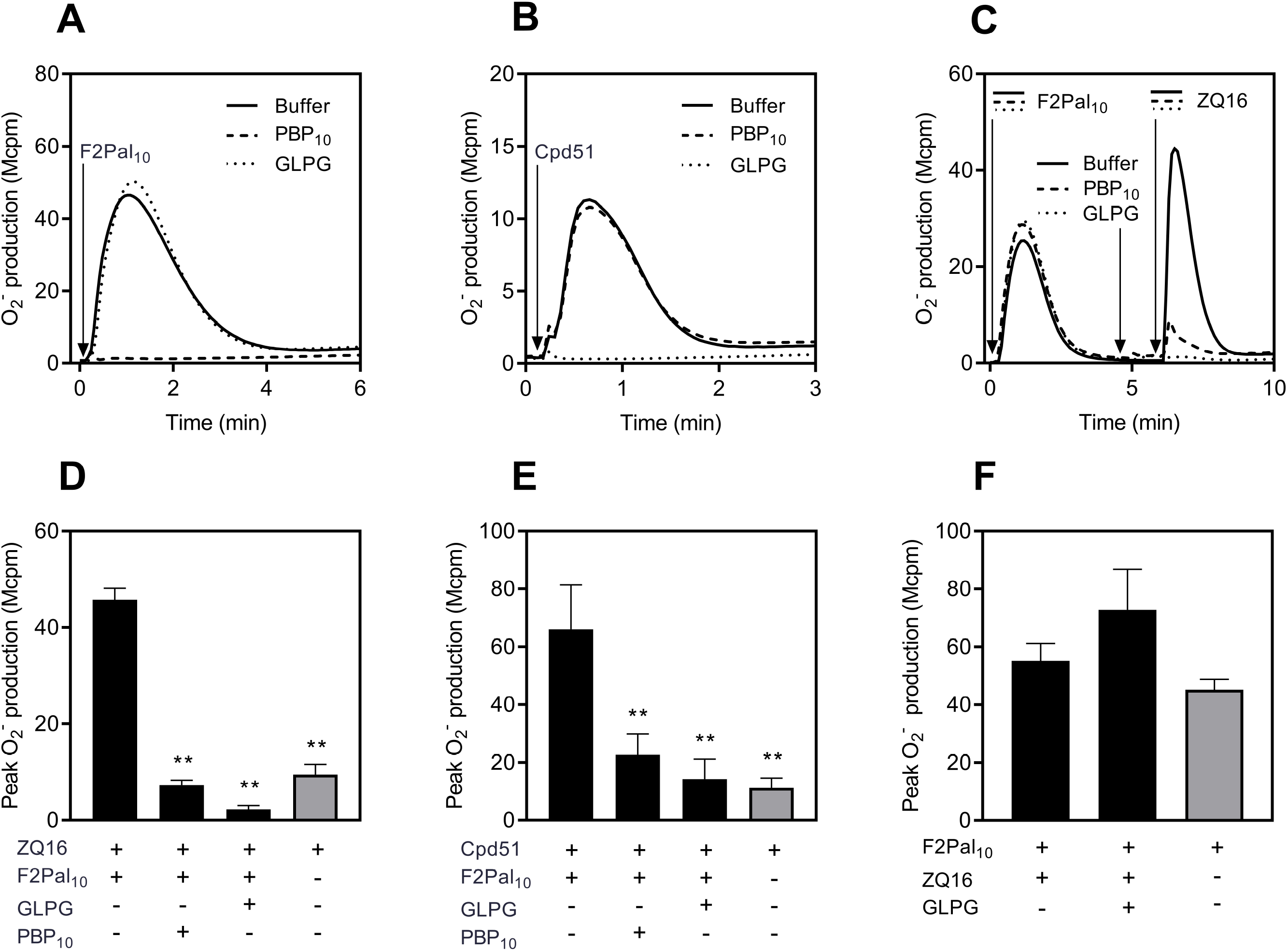
GPR84 agonists trigger a unidirectional reactivation of F2Pal_10_ desensitized neutrophils. **A-B**) Naïve neutrophils were activated by **A**) F2Pal_10_ (500 nM) or **B**) Cpd51 (10 nM) in the absence (solid line) or presence of the FPR2 specific inhibitor PBP10 (1 μM, dashed line) or the GPR84 specific antagonist GLPG1205 (1 μM, dotted line) and the release of superoxide anions was measured continuously over time. One representative experiment out of three individual experiments for each agonist/antagonist combination is shown. **C**) Naïve neutrophils were first activated with F2Pal_10_ (500 nM, first arrow). When the response had declined, cells received a second treatment with either the FPR2 specific inhibitor PBP10 (1 μM, dashed line), the GPR84 inhibitor GLPG1205 (1 μM, dotted line), or buffer (solid line) for one minute (time of addition is indicated by second arrow), followed by a second stimulation with ZQ16 (1 μM, time of addition is indicated by the third arrow). One representative experiment out of three individual experiments for each agonist/antagonist combination is shown. **D-E**) Summary of the inhibitory effects by receptor specific antagonists on the second response induced by **D**) ZQ16 (1 μM) and **E**) the Cpd51 (100 nM) from F2Pal_10_ desensitized neutrophils (n = 3 - 5). The naïve ZQ16- or Cpd51-response from neutrophils stimulated in the absence of F2Pal_10_ and inhibitors is included for comparison (grey bars). Data are expressed as peak superoxide production (mean + SEM) and statistically significant differences were assessed using a repeated measurement one-way ANOVA followed by Dunnett’s multiple comparison test to the GPR84-agonist induced response in F2Pal_10_ desensitized cells in the absence of inhibitors. **F**) Reversed order of receptor agonist addition, i.e., naïve neutrophils were first activated with buffer or the GPR84 agonist ZQ16 (1 μM). Once the ZQ16-response had declined to base-level, the cells received a second stimulation with the FPR2 agonist F2Pal_10_ (500 nM). The GPR84 antagonist GLPG1205 (1 μM) was added one min before the second F2Pal_10_ stimulation. The bar graph shows the peak of the second F2Pal_10_-induced superoxide response (mean + SEM, n = 3). Statistically significant differences were assessed using a repeated measurement one-way ANOVA followed by Dunnett’s multiple comparison test to the F2Pal_10_ induced response in ZQ16-desensitized cells in the absence of GLPG1205.

### The FPR2 reactivation signals are sensitive to Calyculin A and bypass β-arrestin recruitment

Recent research show that the signals generated by agonist occupied GPCRs differ depending on the agonist that initiates signaling, of which some, in addition to activating the heterotrimeric G protein, also may recruit β-arrestin to initiate non-canonical signaling (33). The FPR2 agonist F2Pal_10_ activates a Gαi containing G-proteins but lacks the ability to recruit β-arrestin, and the signals downstream the activated receptor is thus biased (18, 34, 35). The data obtained using F2Pal_10_ as an FPR2 desensitizing agent clearly show that β-arrestin recruitment downstream FPR2 is not a feature that is required for the amplification the of GPR84 response.

To investigate the importance of β-arrestin recruitment for the ability of GPR84 mediated receptor cross-talk with desensitized FPR2s, we determined the cross talk activation induced by DL175, a compound recognized by GPR84 and recently demonstrated to lack the ability to recruit β-arrestin (25). We could confirm the described GPR84 signaling property of DL175 as demonstrated by a strong signaling bias towards the G-protein-cAMP pathway (signaling achieved without any β-arrestin recruitment) also with high concentrations (100 μM) of DL175 (Fig 5A, B). The signaling pattern of DL175 was opposite to the signaling characteristics of ZQ16 that triggers GPR84 down-stream activation of the Gαi signaling pathway as well as recruitment of β-arrestin with EC50 values of ~ 5 nM and ~ 3 μM, respectively (Fig 5A, B). Despite the signaling bias of DL175 away from β-arrestin recruitment, the DL175-induced ROS response was robustly amplified in FPR2-desensitized neutrophils (Fig 5C). Further, also this DL175 response was inhibited not only by the GPR84 antagonist but also by the FPR2 antagonist PBP10, and the level of response reached in the presence of PBP10 was comparable to that induced by DL175 in naïve neutrophils (Fig 5D). Thus, the data obtained with DL175 show that although this GPR84 agonist is unable to recruit β-arrestin down-stream of GPR84, the signals generated reactivate desensitized FPR2. Together with the fact that also F2Pal_10_ lacks ability to recruit β-arrestin, we conclude that β-arrestin does not play a role in the receptor cross-talk induced FPR2-reactivation process.

**Figure 5.**
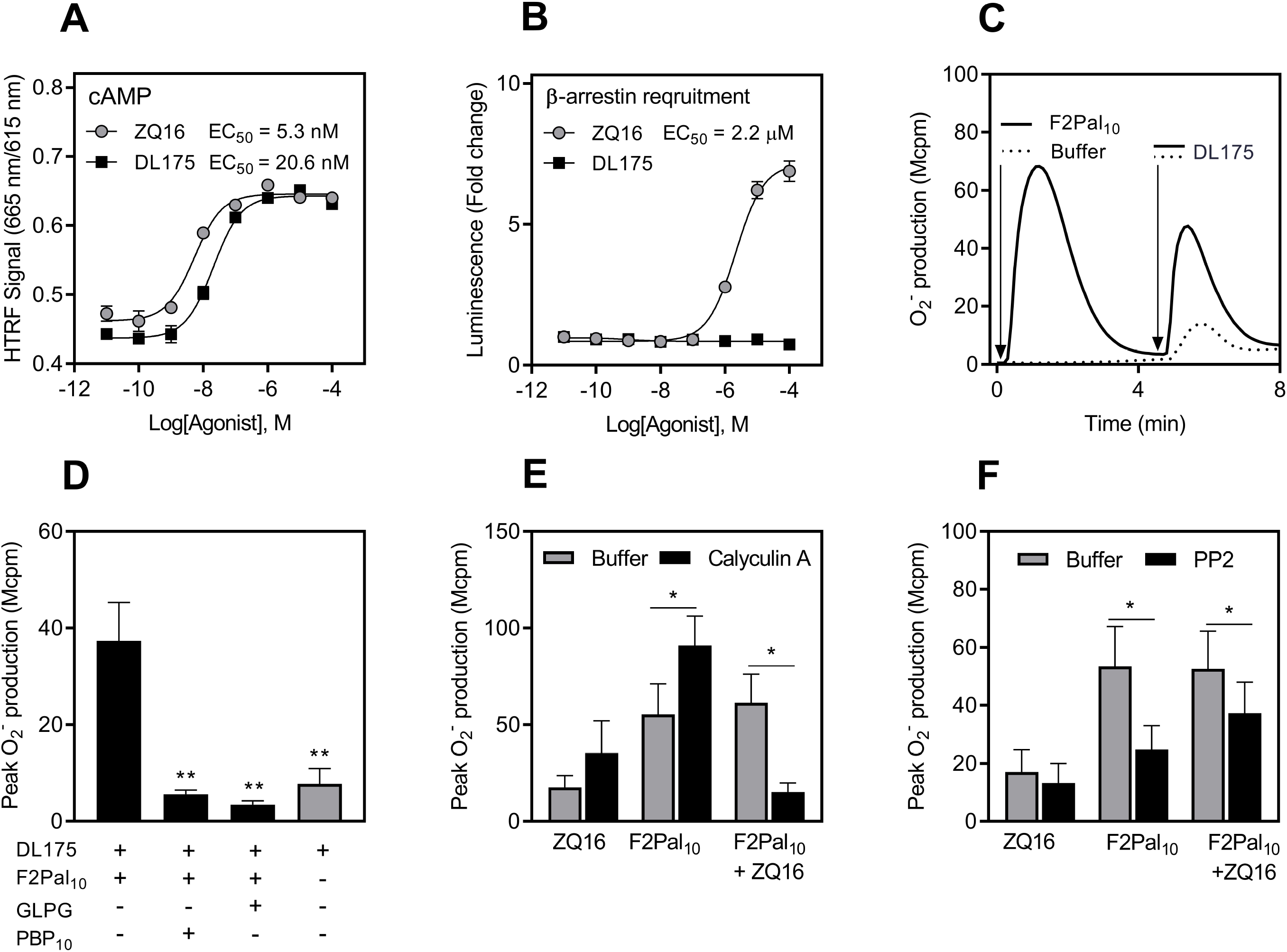
FPR2 reactivation through GPR84 occurs independent of GPR84-induced β-arrestin recruitment and involves Calyculin A sensitive phosphatases. **A-B**) GPR84 overexpressing HEK 293 cells were activated with different concentrations of ZQ16 or DL175. Dose-response curves of agonist-induced **A**) cAMP or **B**) recruitment of β-arrestin was measured. The EC50 values were calculated (mean + SEM, n = 3 independent experiments). **C**) Neutrophils were first activated with F2Pal_10_ (500 nM, solid line) or buffer (dotted line) and the release of superoxide anions was measured continuously over time. After five min, when the F2Pal_10_ response had returned to basal levels, the neutrophils received a second stimulation with DL175 (5 μM). One representative experiment out of four individual experiments is shown. **D**) The bar graph shows summary of the second peak DL175-induced superoxide response from F2Pal_10_-desensitized neutrophils treated for one minute with or without GLPG1205 (1 μM) or PBP10 (1 μM) prior DL175 stimulation (mean + SEM, n = 4). The naïve DL175 response from neutrophils stimulated in the absence of F2Pal_10_ and inhibitors is included for comparison (grey bar). Statistically significant differences were assessed using a repeated measurement one-way ANOVA followed by Dunnett’s multiple comparison test to the DL175 response in F2Pal_10_ desensitized cells in the absence of inhibitors. **E-F**) Neutrophils pre-incubated with buffer (grey bars), **E**) Calyculin A (60 nM, 10 min, black bars) or **F**) PP2 (500 nM, 10 min, black bars) were activated with F2Pal_10_ (500 nM) or buffer (naïve cells) before triggered by ZQ16 (1 μM) or F2Pal_10_ (500 nM). The bar graphs (mean + SEM, n = 4) show the peak superoxide release induced by F2Pal_10_ and ZQ16 in naïve neutrophils as well as by ZQ16 in F2Pal_10_-desensitized neutrophils, pre-treated in the absence (grey bars) and presence of Calyculin A or PP2 (black bars). The effects of pharmacological inhibitors in comparison to buffer treated cells for each agonist combination (buffer/ZQ16, buffer/ F2Pal_10_ and F2Pal_10_/ ZQ16) were analyzed by a paired Student’s *t*-test.

To gain further insights into the intracellular signals involved in the receptor cross-talk transduced by GPR84 and leading to reactivation of the desensitized FPR2, the effects of two pharmacological inhibitors that potently inhibit PP2A/PP1 phosphatases (Calyculin A) and Src kinases (PP2), respectively, were determined. F2Pal_10_-desensitized neutrophils were treated with specific pharmacological inhibitors followed by FPR2 reactivation with the GPR84 agonist ZQ16. The presence of Calyculin A significantly reduced the cross-talk induced ROS production (Fig 5E). Of note, this was not only due to an inhibition on the naïve GPR84- or FPR2-response, as the effect of Calyculin A, if any, on the ZQ16-and F2Pal_10_-induced ROS production in naïve non-desensitized neutrophils was increased rather than reduced (Fig 5E). The Src kinase inhibitor PP2 reduced the naïve F2Pal_10_ response as well as the receptor cross-talk induced ROS production (Fig 5F).

Taken together, these data show that, at the signaling level, β-arrestin is not involved in the GPR84-mediated FPR2 reactivation, a process that engages Calyculin A sensitive phosphatases.

### ROS production induced by GPR84 agonists in FPR1-desensitized neutrophils

Previous studies of FPR-signaling (summarized in some recent reviews (4, 36) have shown that the two neutrophil FPRs (FPR1 and FPR2) regulate the activity of the neutrophil NADPH-oxidase in almost identical manners (37). In accordance with this, desensitized FPR1 and FPR2 both are reactivated by signals generated by the agonist occupied PAFR (shown for FPR1 in Fig 6A) and the primed PAF response is inhibited not only by a PAFR antagonist but also by a FPR specific antagonist (38). Thus, a process of receptor cross-talk induced receptor reactivation is disclosed for both FPR1 and FPR2 (4). The basic characteristics of this process is shared by the GPR84 agonist triggered activation of FPR2-desensitized neutrophils (Fig 2 and Fig 4). Based on these facts, we assumed that also the desensitized FPR1 should be reactivated by agonist-occupied GPR84 and that the receptor cross-talk should result in an amplification of the neutrophil response as a result of FPR1 reactivation. Using the FPR2 desensitization and GPR84 agonist-induced reactivation protocol but with the FPR2 agonist replaced by the potent FPR1 specific agonist fMIFL, we found, however, that the response by GPR84 agonists induced in FPR1 desensitized neutrophils was unaffected or even reduced (Fig 6B, C). We intentionally used low nanomolar range concentrations (0.1-10 nM) of fMIFL as this peptide is a full agonist with an EC50 of ~ 0.1 nM (39). Similar heterologous receptor cross-talk activation profile obtained with FPR1 desensitized neutrophils and ZQ16 was obtained when this GPR84 agonist was replaced with the other GPR84 agonist Cpd51 (Fig 6D), and when the FPR1 agonist fMIFL was replaced with the most commonly used FPR1 agonist fMLF as desensitizing agent (Fig 6E).

**Figure 6.**
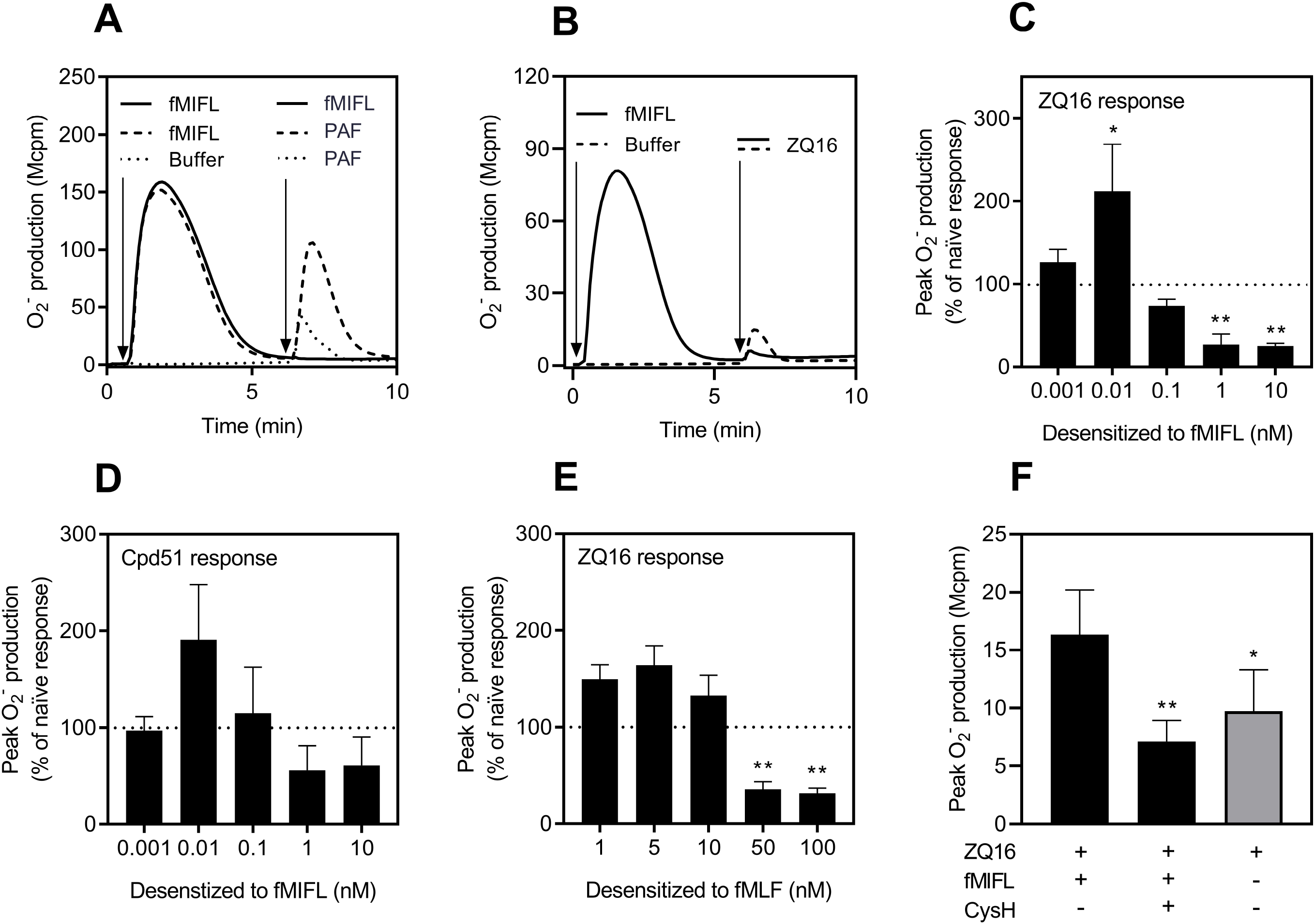
Cross-talk between FPR1 and GPR84. **A**) Neutrophils were first stimulated with fMIFL (10 nM, solid and dashed lines, time of addition indicated by the first arrow) or buffer (dotted line). When the responses had returned to basal levels, the cells received a second stimulation (time of addition indicated by the second arrow) with fMIFL (10 nM, solid line) or PAF (100 nM, dashed and dotted lines). One representative experiment is shown. **B**) Neutrophils were activated with fMIFL (10 nM, solid line, time of addition indicated by the first arrow) or buffer (dashed line). Once the fMIFL response had returned to base-level, the cells received a second stimulation with ZQ16 (1 uM, time of addition indicated with the second arrow). One representative experiment out of four individual experiments for each agonist combination is shown. **C-E**) The bar graphs (mean + SEM, n = 5 - 9) show a summary of the second ROS response induced by **C**) ZQ16 (1 μM) or **D**) Cpd51 (1 nM) in neutrophils pre-activated/desensitized with different concentrations fMIFL and **E**) induced by ZQ16 (1 μM) in neutrophils desensitized with different concentrations of fMLF. The data is expressed as % of the naïve GPR84 agonist response (the dotted lines indicate the 100% of naïve response, peak superoxide production induced by GPR84 agonists in the absence of FPR1 agonists). To adjust for the increased background levels after FPR1 stimulation, the baseline activity just before the second stimulation was subtracted from the second GPR84-agonist induced peak superoxide responses. Statistically significant differences were assessed using a repeated measurement one-way ANOVA followed by Dunnett’s multiple comparison to the naïve GPR84 response. **F)** Neutrophils were stimulated with buffer (naïve neutrophils) or pre-activated/desensitized with fMIFL (0.01 nM). Once the fMIFL-induced superoxide release had declined, the cells received a second treatment with buffer or the FPR1 antagonist Cyclosporin H (CysH, 1 μM) for one minute followed by a second stimulation with ZQ16 (1 μM). The bar graph shows the second peak superoxide response (mean + SEM, n = 3) with the naïve ZQ16 response (grey bar) included for comparison. Statistically significant differences were assessed using a repeated measurement one-way ANOVA followed by Dunnett’s multiple comparison to the ZQ16-induced response in fMIFL-desensitized cells in the absence of CysH.

The responses induced by GPR84 agonists in FPR desensitized neutrophils highlight a fundamental difference between the two FPRs in relation to GPR84, a difference that is clearly seen when the concentrations of the FPR agonists used, generate a full or near full neutrophil response. As shown in Fig 2, the GPR84-triggered response in FPR2-desensitized neutrophils was as pronounced or even higher (for ZQ16) than that induced by F2Pal_10_ in naïve neutrophils. In contrast, the GPR84-triggered response in neutrophils desensitized with corresponding high concentrations of an FPR1 agonist was not increased but rather suppressed (Fig 6B-E). It should be noticed, however, that when the desensitizing FPR1 agonist concentration was reduced, the GPR84 induced response was somewhat potentiated (Fig 6C-E). The modest potentiation of the ZQ16 response in fMIFL (10 pM) desensitized cells was sensitive to the FPR1 antagonist Cyclosporine H (Fig 6F), suggesting that a reactivation of the desensitized FPR1 is achieved when the desensitizing agonist fMIFL concentration is too low to heterologously desensitize GPR84.

Taken together, our data clearly show that both FPR2 and FPR1 can communicate with GPR84 to regulate ROS production in human neutrophils. In addition, it is obvious that whereas the GPR84 triggered ROS production is amplified in FPR2 desensitized neutrophils, FPR1 agonists primarily desensitize GPR84.

## Discussion

Agonists recognized by the medium-chain fatty acid receptor GPR84 are fairly poor activators of the ROS generating NADPH-oxidase in naïve neutrophils, but the agonists are transferred to potent activator of the oxidase in FPR2 (formyl peptide receptor 2) desensitized neutrophils. The augmentation of the GPR84 triggered response is achieved through an earlier described receptor cross-talk mechanism in neutrophils (4, 17), initiated by receptor down-stream signals generated by GPR84, that reactivate the desensitized FPR2. Neutrophils express in addition to FPR2, the closely related FPR1 and although both receptors recognize *N*-formylated peptides, their respective orthosteric agonist binding pocket differ and by that they have different agonist recognition profiles (30). In accordance with the large similarities between these receptors in their cytoplasmic domains (36), signaling by the two receptors in particular regarding neutrophil ROS production have in our earlier studies been shown to be very similar or even qualitatively identical (37). This study reveals, however, a fundamental difference between FPR1 and FPR2 in their respective communication with GPR84. An in-depth characterization of the receptor cross-talk patterns shows that GPR84 utilizes signaling through a pathway that triggers a reactivation of desensitized FPR2 and by that, GPR84 agonists independent of their ability to recruit β-arrestin are transferred to potent inducers of ROS production. Despite the fact that the signals generated by GPR84 can reactivate also FPR1-desensitized neutrophils, this receptor primarily suppress ROS production induced by GPR84 agonists. As the FPRs as well as GPR84 are emerging as promising targets for treating inflammation-associated diseases (23, 40), the novel receptor cross-talk between the two FPRs and GPR84 described, will increase the understanding of the complex signaling network that regulates GPCR-mediated neutrophil activation and the inflammatory response.

The neutrophil production/release of ROS induced by FPR agonists follow a typical GPCR-mediated response pattern in neutrophils, i.e., the time for an active ROS production/release is rather short and the response is after a period of minutes fairly rapidly terminated through homologous receptor desensitization (41). Accordingly, the homologous FPR-desensitized neutrophils are no longer responsive to an activation by the same agonist or to another agonist that binds to the same receptor. FPR-desensitized neutrophils produce/release, however, ROS when activated by an agonist that are recognized by the receptor for platelet activating factor (PAFR). In fact, PAF is a poor activator of naïve neutrophils, but this agonist is turned into a potent activator of the ROS producing system in FPR1 as well as FPR2 desensitized neutrophils (17, 38). It is clear that the level of ROS produced by naïve neutrophils when activated by specific GPR84 agonists (both ZQ16 and the structurally related analog Cpd51) is fairly low and almost negligible as compared to the ROS release following neutrophil activation with agonists specifically recognized by FPR1 or FPR2. Interestingly, we show that the response induced by GPR84 agonists share the same activation profile as PAF in relation to FPR2-desensitized neutrophils, characterized by an amplification of the response that reach levels comparable to that induced by FPR agonists in naïve neutrophils. The fact that this amplification applied to several different FPR2-as well as GPR84-specific agonists suggests that the effect lies at the receptor level and is independent of the particular agonist pair used to desensitize FPR2 and activate GPR84, respectively. Although it is known that FPR2 has the capacity to regulate the functions of many other GPCRs in human neutrophils (4) the interplay between FPR2 and GPR84 has previously not been studied. Thus, this study does not only put GPR84 on the list of GPCRs that can cross-talk with the FPRs but also further support the emerging notion that GPCR activity is regulated at multiples levels through complex mechanisms.

We have earlier described the regulatory role of FPRs and the position they have in the receptor hierarchy to the other neutrophil receptors such as the PAFR and the ATP receptor P2Y2R (4, 16–18, 30, 38). Based on the results presented in these studies we have put forward a receptor signaling cross-talk activation model for reactivation of desensitized FPRs; reactivation of the FPRs is initiated by agonist binding to the PAFR/P2Y2R, and the signals generated down-stream of the G-protein activated by these receptors, activate the desensitized FPRs from the cytosolic side of the membrane (4). Even though the precise signals leading to an amplified GPR84 response in FPR2-desensitized neutrophils remains to be elucidated, the results obtained with these two receptors together fit perfectly to our receptor signaling cross-talk activation model for receptor reactivation. Further, it is clear from the data presented that the reactivation signals utilized in the described cross-talk are not the same as those used by the FPRs to directly activate the NADPH-oxidase. This is illustrated by the facts that Calyculin A, a specific phosphatase inhibitor earlier shown to potentiate the response induced in naïve neutrophils, blocks the cross-talk signaling leading to reactivation of FPR2-desensitized neutrophils not only when PAF is used to activate the neutrophils (17), but also with the reactivation is induced by GPR84 agonists. To further identify signaling pathways of importance for the GPR84 triggered reactivation of desensitized FPR2, we included a biased GPR84 agonist (DL175) unable to trigger recruitment of β-arrestin. The data obtained DL175 showed that the receptor down-stream signals generated by GPR84 reactivate FPR2 desensitized neutrophils also with this agonist which suggests that the reactivating signal(s) are generated independent of β-arrestin recruitment. The fact that also neutrophils activated/desensitized with FPR2 agonists unable to recruit β-arrestin (e.g., F2Pal_10_ and PSMα2) are reactivated by the receptor cross-talk signals generated by GPR84 (this study) as well as by the PAFR and P2Y2R (16–18, 35), show that β-arrestin is not at all involved in the receptor cross-talk reactivation of FPR2. It would be highly interesting to explore the effect on neutrophil activation of a biased GPR84 agonist that is in favor of β-arrestin recruitment but to our knowledge, such a biased agonist has not yet been identified.

Neutrophils express two structurally very similar FPRs (FPR1 and FPR2), and the two receptors also share many similarities regarding the transduced receptor down-stream signals as well as the cellular responses (30, 36). Earlier data obtained in studies designed to determine receptor hierarchies, suggest that FPR1 has the same position as FPR2 in the neutrophil GPCR hierarchy. This suggestion is, however, challenged by the results presented in this study. It is clear that FPR1-desensitized neutrophils similar to FPR2-desensitized neutrophils can be reactivated by the receptor cross-talk reactivation signals generated by GPR84 giving rise to an amplified ROS-response. However, in contrast to FPR2, this can only be achieved with low concentrations of the FPR1-agonist (~ 0.01 nM and ~ 5 nM for fMIFL and fMLF, respectively), well below the EC50 concentrations for activation with these agonists (0.1 nM and 50 nM for fMIFL and fMLF, respectively, (39)). In contrast, the very low response induced by GPR84 agonists in naïve neutrophils is not affected or even reduced when the desensitized state is induced by higher concentrations of the FPR1-agonists. The mechanisms that regulate the functional duality of the FPR1 agonists, having the capacity to reduce the activity induced by GPR84 agonists on the one hand and at the other hand allow a reactivation of FPR1 desensitized neutrophils, remains to be elucidated. Speculatively, one possibility is that the desensitized state of both FPR1 and FPR2 are transferred to active signaling states provided that the receptor reactivation signals are generated. As such, no signal would be generated by GPR84 when high concentrations of the FPR1 agonist have been used to desensitize the receptor. That would mean that the non-responding heterologous desensitized state of GPR84 is generated by high concentrations of FPR1 agonists whereas FPR2 lacks the ability to desensitize GPR84 irrespectively of concentration. The non-responding heterologous desensitized state of GPR84, induced by higher concentrations of FPR1 agonists, places GPR84 in the same hierarchy category as CXCR1/2 (receptor for IL-8) that has been shown to be regulated in a similar fashion. That is, so called endpoint chemoattractants specific for FPR1 heterologously desensitize the neutrophil response to agonist recognized by CXCR1/2 (12–15). The fact that GPR84 generates receptor cross-talk signals that reactivate FPR1 neutrophils desensitized with lower concentrations of FPR1 agonists, suggests that no heterologous GPR84 desensitization is induced by such concentrations of the FPR1 agonists, whereas the ability to be reactivated remains. The outcome of the GPR84 triggered response in FPR1-desensitized neutrophils would then be dependent on whether the FPR1-agonist concentration used to desensitize FPR1 is sufficient to also desensitize GPR84 or not. The distinct different receptor cross-talk patterns between FPR1 and FPR2 further highlights the emerging novel aspects of the two FPRs in modulating signaling and inflammation in different disease conditions (4, 42, 43). Mechanistically the communication between receptors is undoubtedly a highly regulated process that occurs at multiple levels, and no universal mechanism is responsible for the different cross-talk processes. Among many proposed mechanisms and signals involved in receptor cross-talk, receptor cross-phosphorylation by protein kinases such as protein kinase C (PKC) or G-protein coupled receptor kinases (GRKs) that are activated by another receptor upon agonist binding have been suggested to play important roles in heterologous desensitization (44, 45). Future studies should examine the role of different isoforms of GRKs in regulating FPR desensitization and reactivation. Among the known cross-talk patterns, heterologous desensitization of one receptor upon activation of another receptor is more commonly noticed, compared to receptor reactivation by signals generated downstream another receptor which is less frequently observed. More mechanistic studies such as network-based global signaling analysis and advanced experimental designs in primary human cells are highly desired.

In summary, our data further support that receptor cross-talk between different GPCRs represents an important mechanism to control neutrophil function. Our results of the receptor cross-talk signaling patterns mediated by GPR84 specific agonists in FPR desensitized neutrophils reveals a not earlier described signaling difference between FPR1 and the closely related FPR2. Understanding the complex mechanisms underlying receptor communication and intracellular signaling pathways in neutrophils should undoubtedly increase our knowledge of GPCR signaling. In addition, increased knowledge of how such responses are modulated in general would facilitate the development of better GPCR-based functional regulators that can be used to control a wide array of pathologies.

## Abbreviations

cAMP: cyclic adenosine monophosphate
CL: chemiluminescence
CysH: Cyclosporine H
DMEM: Dulbecco’s Modified Eagle Medium
DMSO: dimethyl sulfoxide
fMIFL: *N*-formyl-Met-Ile-Phe-Leu
fMLF: *N*-formyl-Met-Leu-Phe
FPR: formyl peptide receptor
GPCR: G protein-coupled receptor
HRP: horseradish peroxidase
KRG: Krebs-Ringer phosphate buffer
MCFAs: medium chain fatty acids
MCT: mitochondrial cryptic peptide
NADPH: nicotine adenine dinucleotide phosphate
PAF: platelet activating factor
ROS: reactive oxygen species

## Acknowledgement

This work was supported by the Swedish Research Council, the Swedish government under the ALF-agreement, the Elisabeth and Alfred Ahlqvist foundation - Swedish Pharmacy Society grant, the Rune and Ulla Almlövs Foundation grant, the Magnus Bergwall Foundation grant, and the Ingabritt and Arne Lundberg Foundation. Asmita Manandhar is supported by a scholarship from the Novo Nordisk Foundation. We thank senior professor Claes Dahlgren for critical discussions and valuable suggestions during preparation of the manuscript.

## Authors’ contributions

HF, LB, MS, JM designed and oversaw all the aspects of the study. JM, MS, AM, LI, LZ and LB performed and analyzed the experiments with input from TU, XX and HF. HF wrote the first draft and all authors revised the manuscript and approved the final version prior submission.

